# Optimization of AAV6 transduction enhances site-specific genome editing of primary human lymphocytes

**DOI:** 10.1101/2021.05.03.440656

**Authors:** Geoffrey L. Rogers, Chun Huang, Robert Clark, Eduardo Seclen, Hsu-Yu Chen, Paula M. Cannon

**Author notes:** <u>Corresponding Author:</u> Paula Cannon, PhD, Dept. of Molecular Microbiology & Immunology, Keck School of Medicine, University of Southern California, 2011 Zonal Ave, HMR 413A, Los Angeles CA 90033.

## Abstract

Adeno-associated virus serotype 6 (AAV6) is a valuable reagent for genome editing of hematopoietic cells due to its ability to serve as a homology donor template. However, a comprehensive study of AAV6 transduction of hematopoietic cells in culture, with the goal of maximizing *ex vivo* genome editing, has not been reported. Here, we evaluated how the presence of serum, culture volume, transduction time, and electroporation parameters could influence AAV6 transduction. Based on these results, we identified an optimized protocol for genome editing of human lymphocytes based on a short, highly concentrated AAV6 transduction in the absence of serum, followed by electroporation with a targeted nuclease. In human CD4^+^ T cells and B cells, this protocol improved editing rates up to 7-fold and 21-fold respectively, when compared to standard AAV6 transduction protocols described in the literature. As a result, editing frequencies could be maintained using 50-100-fold less AAV6, which also reduced cellular toxicity. Our results highlight the important contribution of cell culture conditions for *ex vivo* genome editing with AAV6 vectors and provide a blueprint for improving AAV6-mediated homology-directed editing of human T and B cells.

## Introduction

*Ex vivo* genome editing of hematopoietic cells has now advanced to the clinic as a treatment for several human genetic and infectious diseases.^1^ The target cells include hematopoietic stem and progenitor cells (HSPCs), capable of reconstituting an entire immune system, as well as more differentiated subsets such as T cells and B cells. One of the most well-studied methods for editing hematopoietic cells combines the transient delivery of a targeted nuclease with transduction of a homology donor DNA template packaged in an adeno-associated virus (AAV) vector.^2–4^ The targeted nuclease is designed for only transient expression, delivered for example by electroporation of zinc-finger nuclease (ZFN) mRNA or Cas9 ribonucleoprotein complexes (RNP). Following introduction of a site-specific break in the targeted chromosomal site, the cellular homology-directed repair (HDR) pathway uses the supplied AAV genome to permanently incorporate modified DNA at that site.^5–7^ These procedures can result in high frequency modification of hematopoietic cells, with editing efficiencies ranging from 20-80% across cell types and genomic loci.^2, 8–20^

In developing protocols for genome editing of hematopoietic cells, significant effort has been expended on the targeted nuclease: developing platforms for efficient transient delivery,^2, 4, 21, 22^ optimizing protein sequences,^23^ and chemically modifying RNA components^24^ to maximize on-target nuclease activity while minimizing potentially deleterious off-target DNA break formation. Additional improvements in HSPC genome editing have focused on identifying culture conditions that facilitate HDR through the manipulation of the cell cycle or DNA repair pathways,^21, 25, 26^ as well as to retain optimal stemness and proliferative potential after engraftment.^8, 11, 12, 21, 27^ In contrast, less attention has been paid to conditions that could affect the delivery of the homology donor DNA using AAV vectors.

The efficacy and safety of AAV for gene delivery is well-studied. The parental virus is a small, nonpathogenic parvovirus, encapsidating a single-stranded DNA genome of about 4.7 kb.^28^ For recombinant vectors, all viral DNA sequences are removed other than the inverted terminal repeats (ITRs) necessary for genome packaging into the capsid. AAV vectors have been used extensively in both preclinical and clinical studies, and two products for the treatment of monogenetic disorders are currently approved by the FDA.^29, 30^ AAV is particularly versatile as a gene therapy vector due to its relatively low immunogenicity, the variety of serotypes available with tropism for different tissues, and the ability to persist as episomal, nonintegrated DNA for upwards of a decade.^31^ As such, most work using AAV has focused on *in vivo* gene delivery, with *ex vivo* applications for AAV comparatively less established.

Screening in human HSPCs,^2, 32, 33^ T cells,^3, 13^ and B cells^16^ identified AAV6 as an effective serotype for transduction *ex vivo* in all three hematopoietic cell lineages. Some recent studies have also reported reagents that may enhance uptake of AAVs, including AAV6, in hepatocytes or human HSPCs.^34, 35^ However, comprehensive work has not been published to optimize the factors that could affect AAV transduction of hematopoietic cells *ex vivo* and thereby influence genome editing outcomes. Here, we report an improved protocol for AAV6 delivery as part of nuclease-mediated genome editing, resulting in improved editing efficiencies in T cells and B cells while also reducing cellular toxicity.

## Results

### Cell culture with serum inhibits AAV transduction

To improve the delivery of homology donor templates based on AAV6 vectors, we first assessed the impact of fetal bovine serum (FBS) concentration in cell culture conditions. Wang *et al*. previously reported that transduction of primary human CD3^+^ T cells with AAV6 in FBS-free conditions improved site-specific genome editing when using multiplicities of infection (MOIs) of 10^4^ – 3 x 10^5^ compared to transduction in media supplemented with 10% FBS.^13^ To determine whether this enhancement was caused by reduced AAV6 transduction in the presence of FBS, we transduced a variety of suspension and adherent cell lines with AAV6 vectors containing a GFP expression cassette (AAV6-CCR5-GFP)^2^ at MOIs of 10^3^ - 10^6^ and quantified any inhibition by 10% FBS (Fig. 1A-B and S1). We observed 77-98% inhibition at the lowest MOIs tested, whereas inhibition was only 0-18% at the highest MOIs, suggesting that the inhibitory factor is dose-limiting and can be out-competed by excess AAV6. Heat inactivation of FBS had no effect on the inhibition of AAV6 transduction, suggesting complement is not involved in this process (Fig. S2).

**Figure 1.**
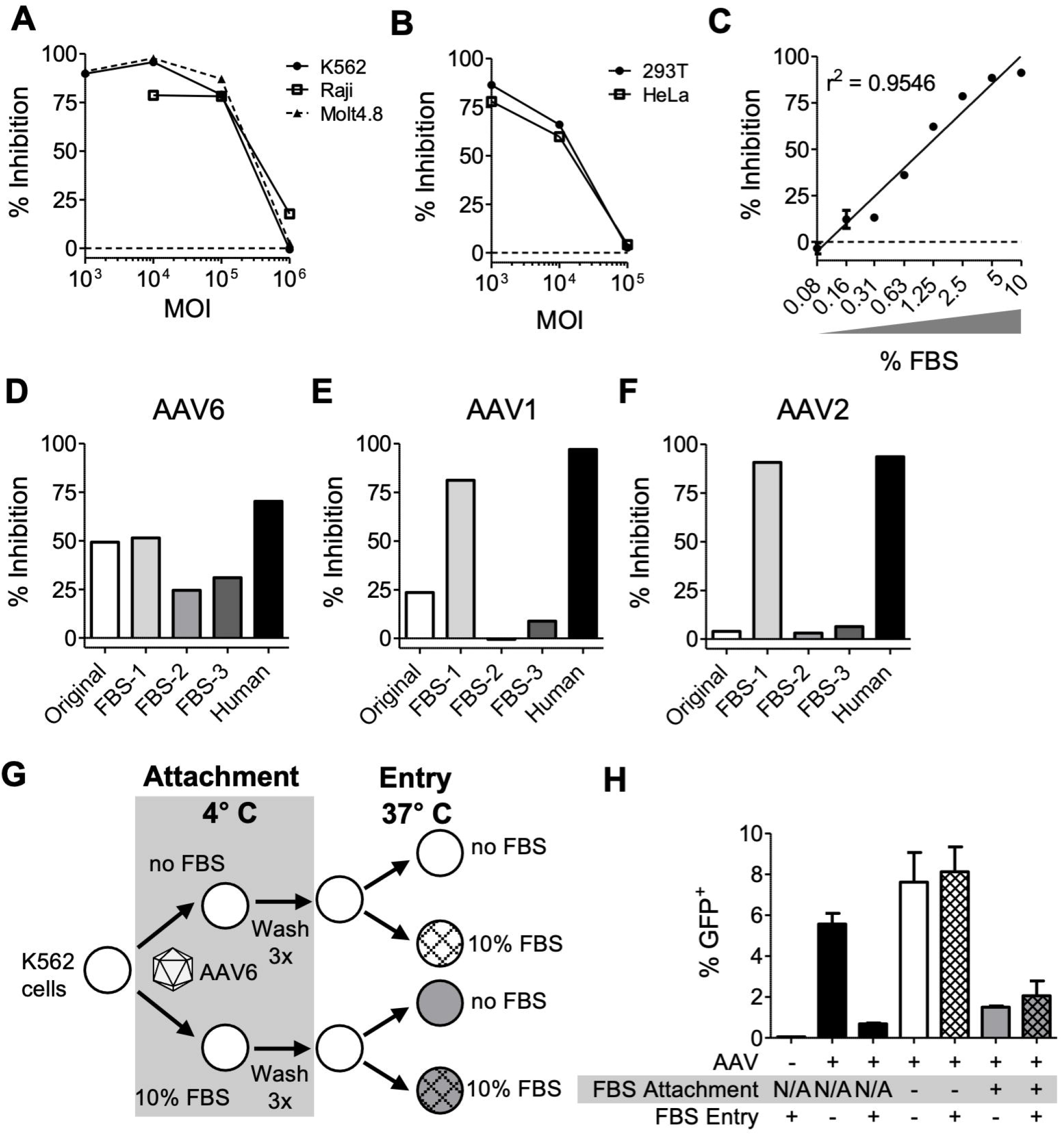
Cell culture serum inhibits AAV transduction. (A) Suspension cell lines were transduced with AAV6-CCR5-GFP vectors at indicated MOIs, with or without 10% FBS, and the percent inhibition compared to no FBS was calculated. Suspension cell lines are shown are K562 (HSPC-like), Raji (B cell), and Molt4.8 (CD4^+^ T cell). Raw data is in Figure S1. (B) Two adherent epithelial cell lines (293T and HeLa) were treated as in (A). (C) Inhibition of AAV6 transduction by MOI = 10^4^ of K562 cells was calculated over a range of FBS concentrations, and a semi-logarithmic regression line was calculated. Data are mean ± SEM for *n* = 3 technical replicates. (D-F) Inhibition of transduction of 293T cells by MOI = 10^4^ of vectors AAV6-CCR5-GFP (D), AAV1-CMV-GFP (E), or AAV2-CMV-GFP (F), by 10% serum for the original batch of FBS used in (A-C), 3 more batches of FBS from different suppliers, and a batch of human AB serum. (G-H) A viral attachment assay was performed for MOI = 10^4^ AAV6-CCR5-GFP vectors on K562 cells, as diagramed (G), and GFP expression was measured after 2 days by flow cytometry (H). Bar colors correspond to treatments, and black bars are control samples transduced at 37°C without prior attachment. Data are mean ± SEM for *n* = 3 technical replicates.

A titration of the FBS concentration during AAV6 transduction of K562 cells at the susceptible MOI of 10^4^ revealed that this inhibitory effect was both potent and dose-dependent (Fig. 1C). Near-complete inhibition was observed at the standard culture concentration of 10% FBS, and some inhibition persisted until FBS was diluted to less than 0.1% of the culture media during transduction.

Next, we investigated the generalizability of this finding across different batches of serum and AAV serotypes using 293T cells, which are permissive to AAV1, AAV2, and AAV6 at an MOI of 10^4^. The original batch of FBS, as well as 3 additional batches, all inhibited AAV6 on 293T cells, ranging between 25-52% inhibition (Fig. 1D). In contrast, significantly more variability was observed with serotypes AAV1 and AAV2, where batch FBS-1 was >75% inhibitory, but FBS-2 and −3 produced minimal inhibition (Fig. 1E-F). Since clinical protocols generally eschew FBS to avoid potential contamination with animal proteins, we also measured the anti-AAV activity of a single batch of human AB serum. This was more strongly inhibitory against all 3 AAV serotypes than any of the batches of FBS (Fig. 1D-F), with the activity against AAV6 exhibiting a similar dose-response curve as was observed with the original batch of FBS (Fig. S3).

Finally, to investigate the mechanism of AAV6 inhibition by FBS, we performed a viral attachment assay. Here, cells are incubated with AAV6 at 4°C to allow viral attachment but not entry, followed by thorough washing and then additional incubation at 37°C to permit uptake of any attached viral particles (Fig. 1G).^36^ As previously observed, AAV6 transduction in control samples maintained throughout at 37°C was inhibited by FBS, and this was also observed when FBS was present during the 4°C attachment step (Fig. 1G). In contrast, allowing AAV6 to first attach to cells at 4°C in the absence of FBS resulted in transduction, regardless of whether FBS was present during the subsequent 37°C incubation. Together, these results suggest that FBS inhibits AAV6 by preventing attachment of the vector to cells rather than acting to prevent viral uptake, or through any changes in cell permissivity at post-entry stages of transduction.

### Impact of culture volume and time on AAV6 transduction

In AAV transduction protocols, MOI is frequently the only characteristic that is reported. However, there are a number of other variables that could impact transduction with the same number of cells and AAV vector genomes. For instance, Ling *et al*. previously reported that increasing cell density during AAV6 exposure improved transduction in both K562 cells and human CD34^+^ HSPCs.^37^ To test this, we transduced equal numbers of K562 cells with AAV6 MOIs ranging from 10^3^ to 10^6^ and using a range of different culture volumes (Fig. 2A). While MOI was clearly an important driver of transduction rates, at each MOI greater than 10^3^, reducing the culture volume also significantly enhanced transduction. These enhancements were sufficient to achieve comparable or superior AAV6 transduction with 10-fold lower MOIs for several comparison points. For example, an MOI of 10^4^ in 5 μL produced 26.6% GFP^+^ cells, but using 500 μL of culture required an MOI of 10^5^ to produce only 20.9% GFP^+^ cells. Interestingly, this impact of culture volume on AAV6 transduction rates was more dramatic at higher MOIs (Table S1), which likely reflects the rules of Brownian motion that govern interactions between viruses and cells in solution.^38^

**Figure 2.**
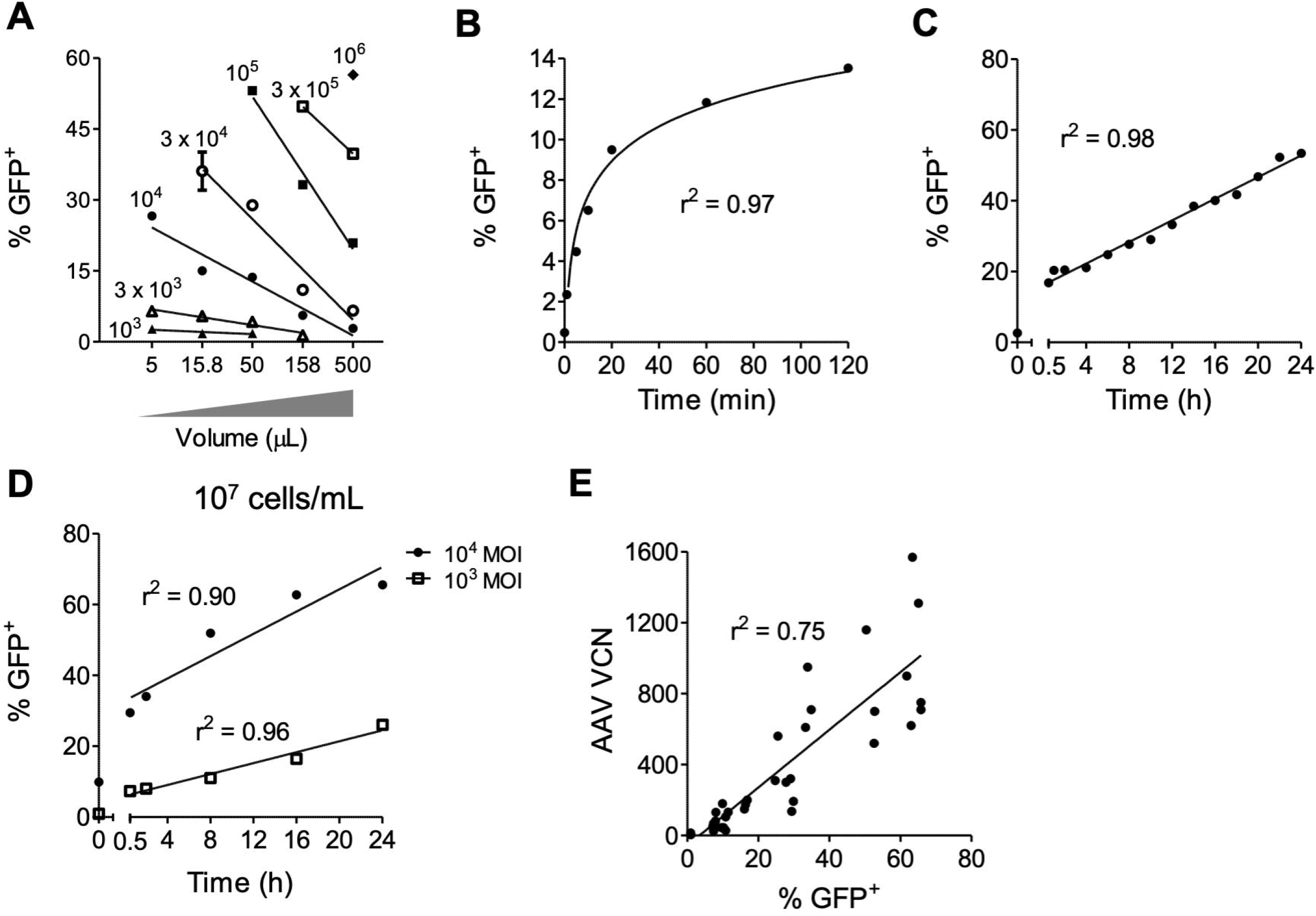
Effects of culture volume and time on AAV6 transduction. (A) The combined influences of AAV MOI and cell culture volume on AAV6 transduction of K562 cells were evaluated. Five x 10^4^ K562 cells were transduced with AAV6-CCR5-GFP at MOIs of 10^3^ – 10^6^ in the indicated volumes for 2 h prior to addition of 10% FBS, and GFP expression was measured after 2 days by flow cytometry. See also Table S1 for regression characteristics. (B-C) K562 cells were transduced with AAV6 at 10^6^ cells/mL and an MOI of 10^4^ for the indicated times. Transduction was halted by 10% FBS, and GFP expression was measured after 2 days. Panel (B) shows that transduction over the first 2 h fits a semi-logarithmic regression, while panel (C) illustrates an initial jump in transduction followed by increases that fit a linear regression starting 0.5 h after transduction. (D) Transduction of K562 cells over time with AAV6 at a higher concentration of 10^7^ cells/mL and MOIs of 10^4^ or 10^3^ was performed as before. See also Table S2 for regression characteristics. (E) Cellular AAV vector copy numbers (VCN) were measured for cells in panel (D) by ddPCR, and a linear regression was used to measure correlation between GFP expression and VCN. Data in panels (A-D) are shown as mean ± SEM for *n* = 3 technical replicates.

As a consideration for AAV6 transduction in the absence of FBS, we were also interested to identify culture times that could optimize transduction while minimizing the deleterious effects of serum starvation on cell viability. Transducing K562 cells at a standard MOI of 10^4^, and in a cell concentration of 10^6^ cells/mL, revealed a biphasic pattern of AAV transduction over time. Specifically, the frequency of GFP^+^ cells increased logarithmically in the first hour or so (Fig. 2B), whereas a linear rate of transduction was observed thereafter up to 24 h of exposure (Fig. 2C). As anticipated, increasing the cell concentration 10-fold to 10^7^ cells/mL while maintaining the MOI of 10^4^ enhanced transduction at all time points tested (compare Fig. 2C and Fig. 2D). Interestingly, regression analysis suggested that the linear rate of transduction was unaltered by the reduced volume, as the slope of the linear regression was not significantly different between the two data sets (Table S2). A lower MOI of 10^3^ significantly altered the slope of the linear regression, but still appeared to display the biphasic rate of transduction (Fig. 2D, Table S2).

Finally, to investigate the physical entry of AAV6 vectors into cells over time, we performed a vector copy number (VCN) analysis on transduced cells. The number of cell-associated AAV genomes showed a strong linear correlation with the percentage of GFP^+^ cells across the different transduction conditions (Fig. 1E), suggesting that the rate of AAV6 transduction over time is regulated by cellular entry of viral particles. Together, these results suggest that the benefits of AAV6 transduction in a low volume of media are realized in the first 1-2 h of exposure, during the logarithmic stage of the biphasic transduction observed.

### Electroporation enhances AAV6 transduction

Electroporation to introduce a targeted nuclease is an additional step in *ex vivo* genome editing protocols that is not performed during AAV transductions. A previous study by Charlesworth *et al*. suggested that recently electroporated cells are more permissive to AAV6 transduction due to a general enhancement of cellular endocytosis.^21^ In agreement, we found that electroporation of K562 cells prior to addition of AAV6 enhanced transduction; however, this was still strongly inhibited by 10% FBS (Fig. 3A). Moreover, this effect was also observed when cells were transduced with AAV6 prior to electroporation (Fig. 3B). FBS was again partially inhibitory, although electroporation rescued transduction to levels comparable to cells transduced without electroporation or FBS. Washing cells to remove AAV6 prior to electroporation did not affect the electroporation enhancement of transduction, suggesting that the AAV6 particles already attached to cells are more efficiently able to transduce cells after electroporation, rather than direct membrane permeation by free AAV. Lastly, we confirmed that electroporation prior to AAV6 exposure was also able to enhance transduction in primary human CD4^+^ T cells and CD34^+^ HSPCs (Fig. 3C-D). Together these results suggest that electroporation enhances AAV transduction of both cell lines and primary human cells, likely through an enhancement of viral uptake, regardless of the sequence of events.

**Figure 3.**
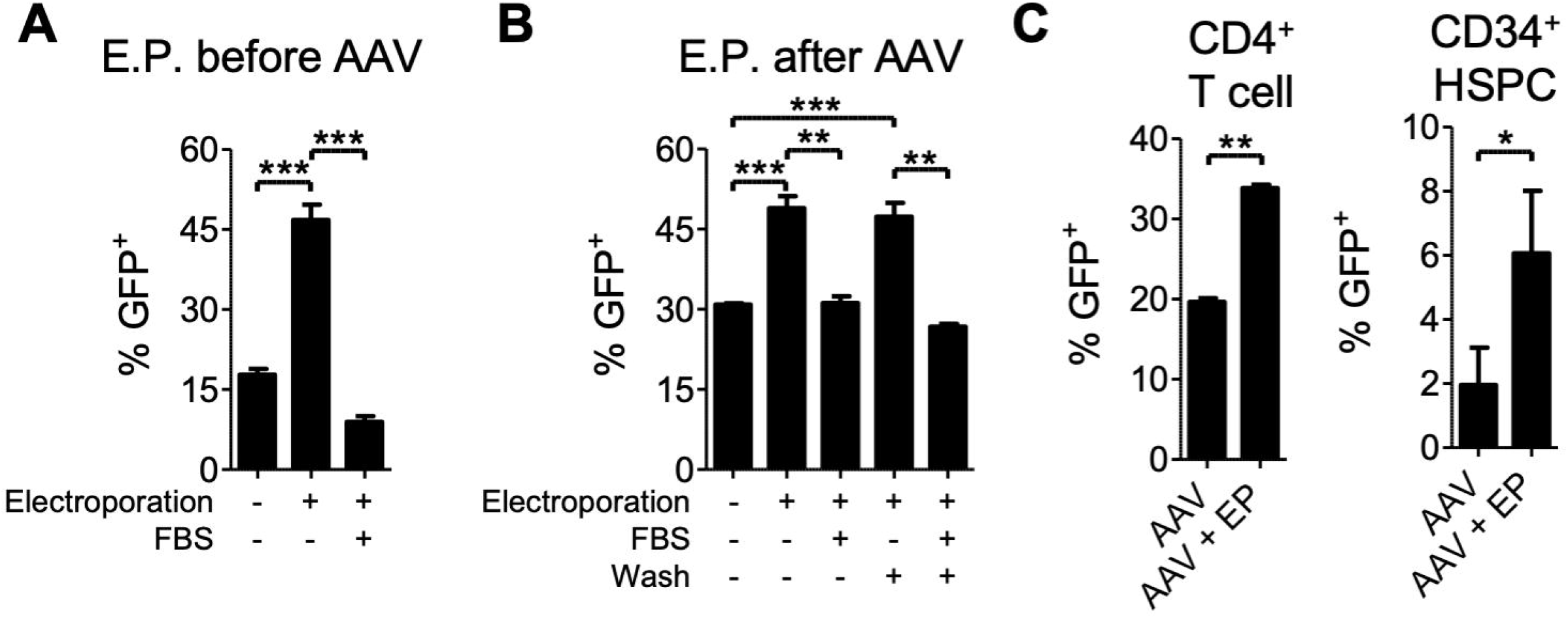
Electroporation enhances transduction by attached AAV6 particles. (A-B) K562 cells were transduced with AAV6-CCR5-GFP vectors at 10^6^ cells/mL and an MOI of 10^4^, and GFP expression was measured after 2 days by flow cytometry for *n* = 3 technical replicates. Electroporation was performed either before (A) or after (B) AAV6 transduction. Washing was performed with PBS after transduction as indicated, and for cells not transduced in the presence of FBS, media was supplemented with 10% FBS after 2 h or after electroporation. (C) CD4^+^ T cells from *n* = 2 human donors were transduced with AAV6-CCR5-GFP at an MOI of 10^4^ with or without prior electroporation, and GFP expression was measured after 2 days by flow cytometry. (D) HSPCs from *n* = 3 donors were transduced with AAV6-CCR5-GFP at an MOI of 3 x 10^3^ with or without prior electroporation, and GFP expression was measured after 1 day by flow cytometry. Data are shown as mean ± SEM. * *p* < 0.05, ** *p* < 0.01, *** *p* < 0.001.

### An optimized AAV6 transduction protocol for nuclease-mediated genome editing

Having identified several parameters that impact AAV6 transduction, we next explored whether optimization of these variables could improve site-specific genome editing by AAV6 homology donors. We used CCR5 editing reagents that are well-validated by our group, combining the AAV6-CCR5-GFP vectors with matched CCR5-specific ZFN mRNAs, to site-specifically insert the GFP expression cassette at CCR5.^2, 39^ We aimed to take advantage of the positive effects we had observed of performing AAV6 transduction in small volumes of media, and the ability of electroporation to enhance AAV6 entry regardless of the relative timing of AAV6 incubation or electroporation. In this way, we designed an optimized protocol where K562 cells were thoroughly washed to remove FBS from culture media, transduced in serum-free media at high concentration of cells (10^7^/mL) using an AAV6 MOI of 10^4^ for 1 hour, and then electroporated with ZFN mRNA and immediately resuspended in media containing 10% FBS (Fig. 4A and Table 1). We contrasted this with a more standard genome editing protocol that involved first electroporating the ZFN mRNA, followed by transduction of cells at a concentration of 10^6^/mL and using an AAV6 MOI of 10^4^ for 2 h in FBS-free media, before the addition of 10% FBS. We observed that the optimized protocol edited 51.1% of cells compared to 39.6% with the original protocol, even though the same amounts of cells, AAV6 vectors and ZFN mRNA were used in each case (Fig. 4B).

**Figure 4.**
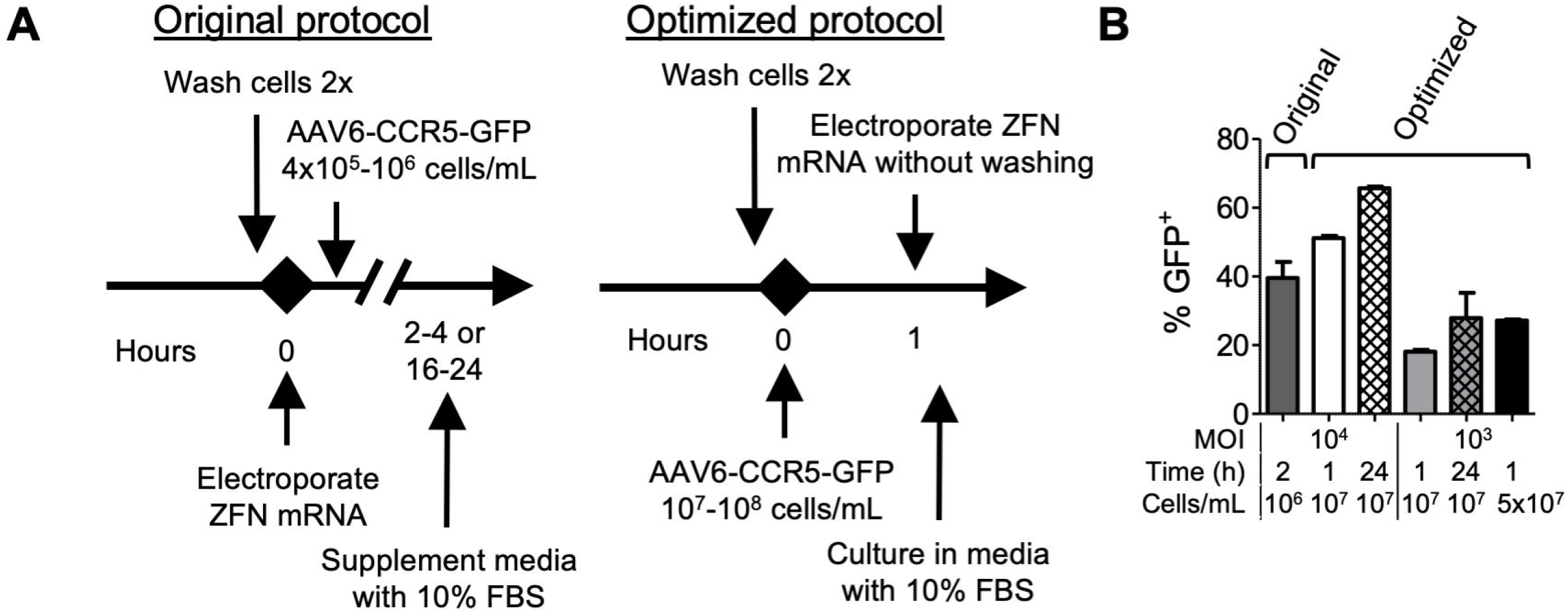
An optimized AAV6 transduction protocol enhances genome editing in K562 cells. (A) Diagram of the original and optimized protocols. In the original protocol, cells are electroporated first, then transduced with AAV6-CCR5-GFP using standard cell culture concentrations prior to addition of FBS. In the optimized protocol, cells are transduced with AAV6-CCR5-GFP at high cell concentrations prior to electroporation, then cultured under standard conditions with FBS supplementation. See Table 1 and Supplementary Methods for additional details of each protocol in specific cell types. (B) K562 cells were genome edited using the original protocol (far left bar) involving AAV6-CCR5-GFP transduction for 2 h after electroporation of ZFN mRNA, or the optimized protocol with varying parameters of MOI, time, and cell concentration as indicated below. Data are shown as mean ± SEM for *n* = 2 technical replicates.

**Table 1.** AAV6 transduction parameters for original and optimized protocols in each cell type. Associated with Fig. 4A.

| **Cell type** | **Transduction Parameter** | **Original Protocol** | **Optimized Protocol** |
| --- | --- | --- | --- |
| **K562 cells** | Concentration (cells/mL) | $10^6$ | $1-5 \times 10^7$ |
|  | Time (hours) | 2 | 1 or 24 |
| **CD4+ T cells** | Concentration (cells/mL) | $10^6$ | $10^8$ |
|  | Time (hours) | 16-24 | 1 |
| **B cells** | Concentration (cells/mL) | $4 \times 10^5$ | $2-5 \times 10^7$ |
|  | Time (hours) | 16-24 | 1 |
| **HSPCs** | Concentration (cells/mL) | $10^6$ | $10^8$ |
|  | Time (hours) | 2-4 | 1 |

We also assessed the impact of the FBS-free transduction period on genome editing rates by transducing the K562 cells for either 1 or 24 h prior to electroporation with ZFN mRNA, and varying the cell concentration. Improvements were observed for the 24 h transduction period at MOIs of both 10^4^ and 10^3^, which could also be achieved by increasing the cell concentration to 5 x 10^7^ cells/mL at the lower MOI with only 1 h incubation (Fig. 4B). However, we consider that a 1 h timeline is likely to be the best choice for primary cells, since this will minimize the duration of serum starvation.

### Improved genome editing in primary human CD4^+^ T cells

We next evaluated genome editing in primary human CD4^+^ T cells, comparing a protocol based on previously published procedures^13, 14^ with our optimized protocol using concentrated AAV6 transduction in FBS-free media for 1 h prior to electroporation (Fig. 4A and Table 1). Across a range of MOIs, the optimized protocol yielded 2.2- to 6.7-fold higher editing levels (Fig. 5A). Significantly, the 33.3% average rate of editing achieved using the optimized protocol at the lowest MOI was greater than the 29.9% editing rate achieved with the previously published protocol at the highest MOI, despite using 50-fold less AAV6. Moreover, no differences were observed in the indel frequencies in cells electroporated with ZFN mRNA alone, suggesting that the optimized protocol did not adversely affect electroporation efficiency or nuclease activity (Fig. 5B).

**Figure 5.**
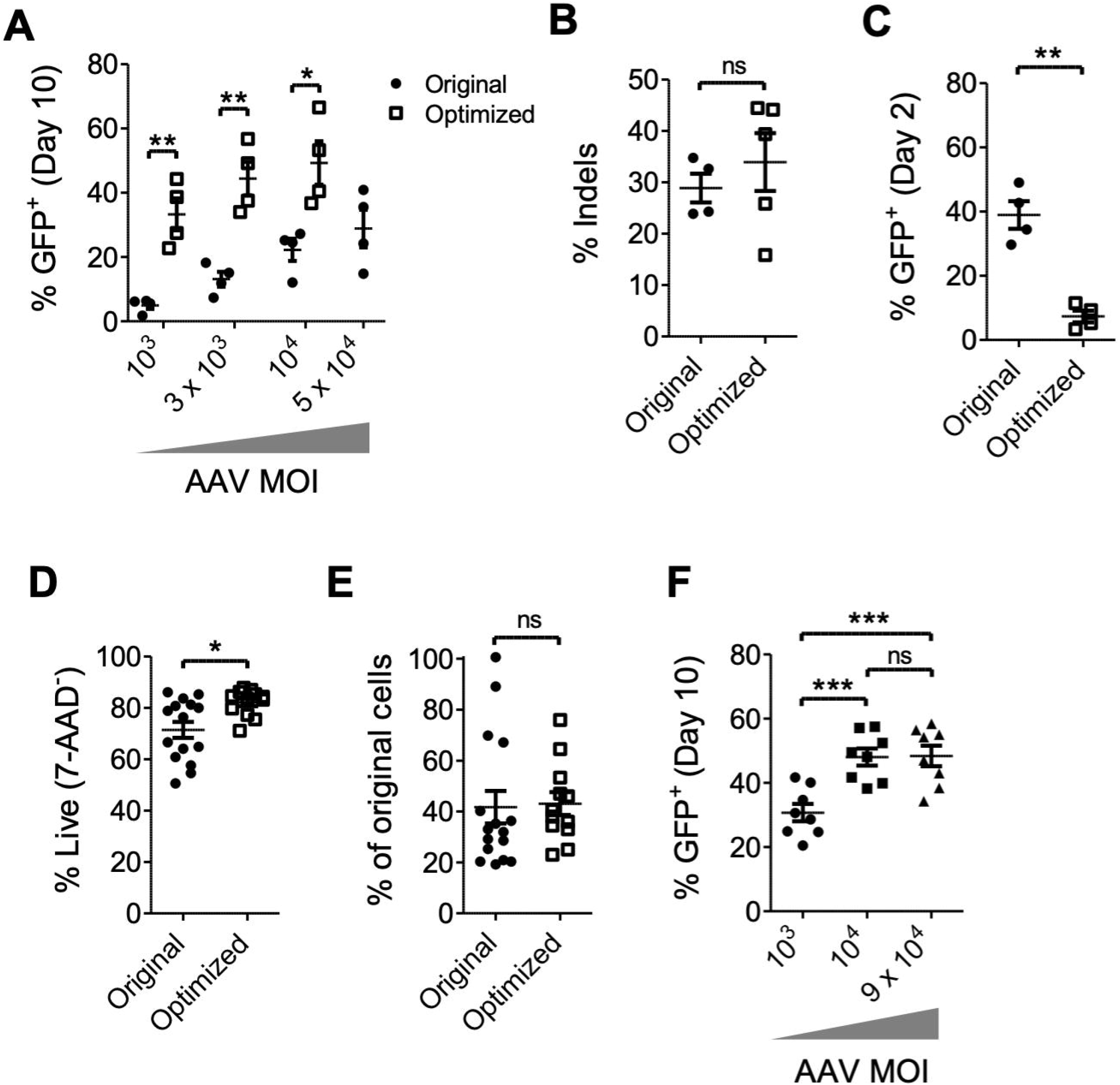
Improved genome editing in CD4^+^ T cells with an optimized protocol. (A-E) CD4^+^ T cells from *n* = 4 human donors were genome edited with either a previously published (original) or optimized protocol at the indicated AAV6-CCR5-GFP MOIs. (A) Stable genome editing shown by GFP expression measured by flow cytometry at day 10. (B) Indel formation at CCR5 by cells electroporated with ZFN mRNA with either protocol. (C) AAV6 transduction with either protocol was measured by flow cytometry for GFP at day 2 in samples treated with AAV6 but not ZFN mRNA. (D-E) Cell viability (D) and total cell counts by hemocytometer (E) were measured 1 day after genome editing. Conditions were pooled across AAV6 MOIs. (F) CD4^+^ T cells from *n* = 8 human donors were genome edited with the optimized protocol at indicated MOIs, and stable genome editing was measured by GFP expression at day 10. Data are shown as mean ± SEM. * *p* < 0.05, ** *p* < 0.01, *** *p* < 0.001, and ns = not significant.

Interestingly, the higher levels of genome editing obtained with the optimized protocol do not appear to have been solely a consequence of improved AAV6 delivery. Indeed, AAV6 transduction in the absence of a targeted nuclease was significantly lower at day 2 using the optimized protocol (Fig. 5C). However, improved cell viability was observed one day after electroporation in cells edited with the optimized protocol, although no significant difference in CD4^+^ T cell numbers was observed at this time point (Fig. 5D-E). It is possible that improved cell health allows higher rates of genome editing, since the DNA repair pathways necessary for HDR are only active during the G2/S phases of the cell cycle.^25, 40^

Finally, we observed a cap on genome editing rates with increasing AAV6 MOIs using the optimized protocol. A 9-fold increase in the AAV6 MOI (from 10^4^ to 9 x 10^4^) with the optimized protocol was unable to increase genome editing rates (Fig. 5F), suggesting that an effective maximum rate had been reached based on additional restrictions on genome editing efficiency.

### Improed genome editing in primary human B cells

We next evaluated whether the optimized AAV6 transduction protocol could improve genome editing rates in primary human B cells, which we have previously found to require AAV6 MOIs of 10^6^ and above for efficient site-specific gene insertion with an original protocol adapted from methods in T lymphocytes^13, 14^ (Fig. 4A and Table 1). Similar to the CD4^+^ T cells, we found that genome editing in primary B cells was greatly enhanced by the optimized protocol, up to 21.1-fold (Fig. 6A). Although the overall maximum rates of editing achieved were the same for both protocols, equal frequencies of GFP^+^ cells were achieved using 10-fold less AAV6 in the optimized protocol, and with only a slight, non-statistically significant reduction observed with 100-fold less AAV6. In contrast to CD4^+^ T cells, transduction of B cells in the absence of a targeted nuclease was slightly improved by the optimized protocol, although this was not statistically significant (Fig. 6B). Lastly, as before, a significant survival advantage was observed with the optimized protocol in terms of B cell viability, as well as the total number of cells remaining one day after electroporation (Fig. 6C-D). Therefore, the optimized protocol allowed for major reductions in the amount of AAV6 vectors required to achieve efficient rates of genome editing in human B cells, although additional barriers beyond AAV6 delivery still appear to limit the maximum genome editing rates that can be achieved.

**Figure 6.**
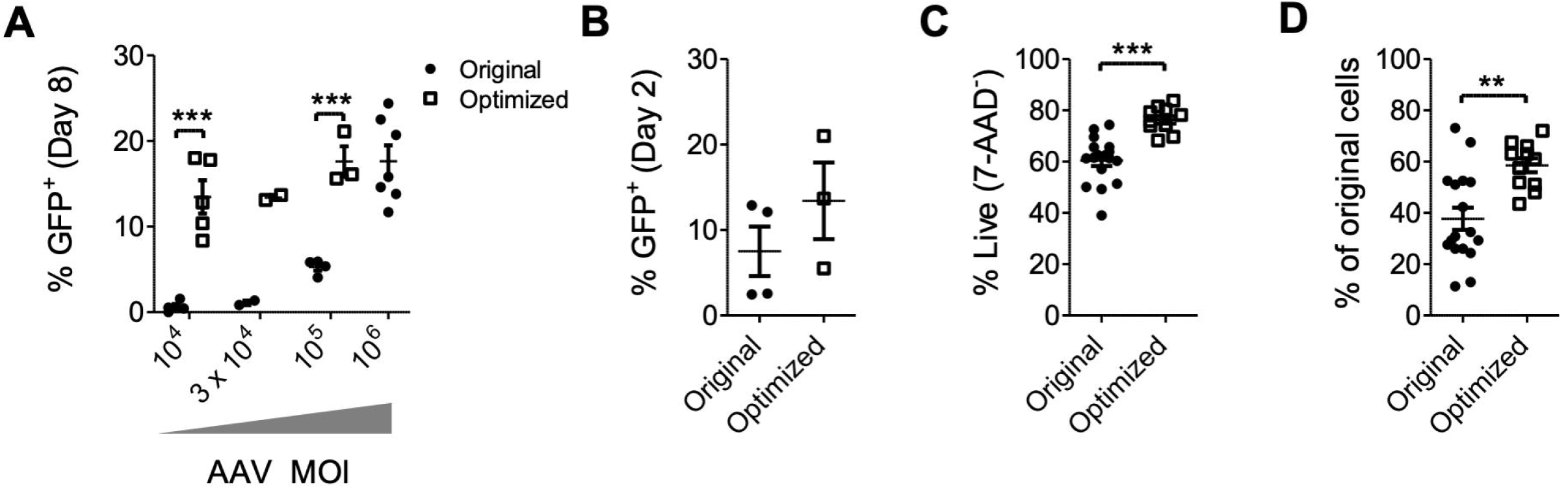
Improved genome editing in B cells with an optimized protocol. CD19^+^ B cells from *n* = 2-6 human donors were edited with either the original or optimized protocols at the indicated AAV6-CCR5-GFP MOIs. (A) Stable genome editing shown by GFP expression measured by flow cytometry at day 8. (B) AAV6 transduction with either protocol was measured by flow cytometry for GFP at day 2 in samples treated with AAV6 but not ZFN mRNA. (C-D) Cell viability (C) and total cell counts by hemocytometer (D) were measured 1 day after genome editing. Conditions were pooled across AAV6 MOIs. Data are shown as mean ± SEM. ** *p* < 0.01, *** *p* < 0.001.

### Genome editing in CD34^+^ HSPCs is unaffected by optimized AAV6 transduction protocol

Finally, we investigated whether our optimized protocol could improve editing in CD34^+^ HSPCs compared to a standardized original protocol used in our lab (Fig. 4A and Table 1). Bulk CD34^+^ HSPCs containing a mixture of stem and progenitor cells have historically required lower AAV6 MOIs for efficient genome editing than human lymphocytes.^2, 13^ However, no differences in editing rates were observed between the protocols across a range of MOIs (Fig. 7A). Transduction in the absence of a targeted nuclease at day 1 was also similar between the 2 protocols (Fig. 7B). There was a slight but significant advantage in cellular viability with the optimized protocol, but no difference in cell numbers 1 day after electroporation (Fig. 7C-D). Thus, despite its impact in primary human lymphocytes, the optimized AAV6 transduction protocol did not appreciably enhance genome editing in CD34^+^ HSPCs.

**Figure 7.**
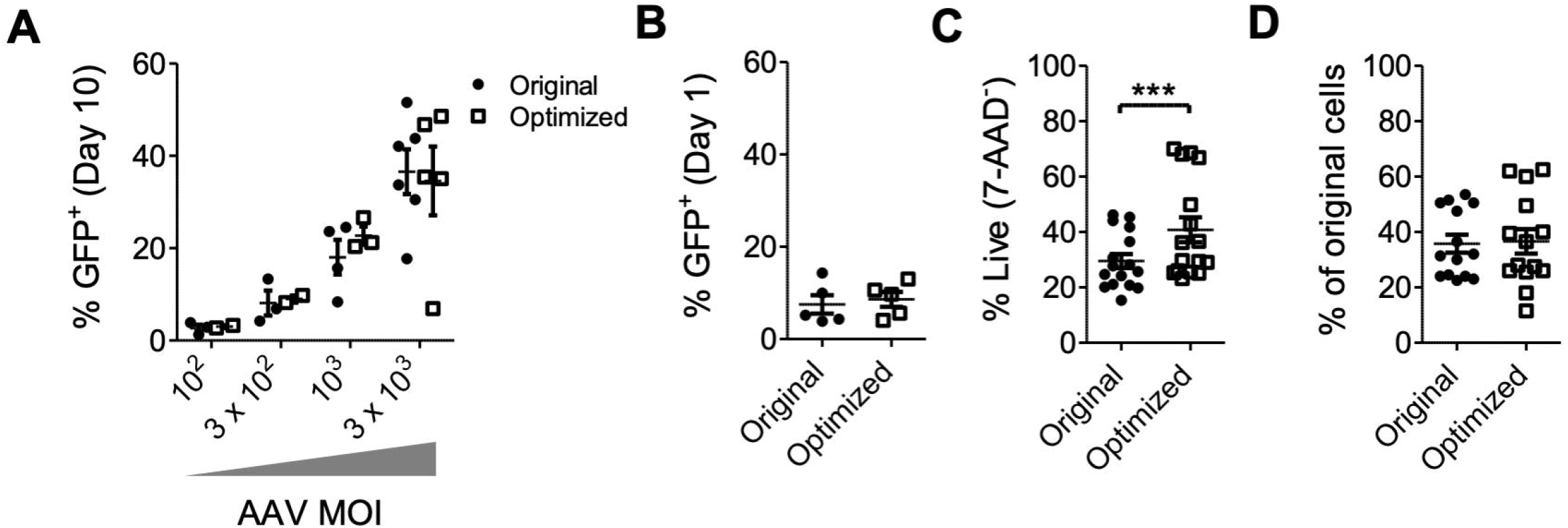
Comparable efficacy of both protocols in HSPCs. CD34+ HSPCs from *n* = 2-6 human donors were edited with either an original or optimized protocol at the indicated AAV6-CCR5-GFP MOIs. (A) Stable genome editing rates, shown by GFP expression measured by flow cytometry at day 10. (B) AAV6 transduction with either protocol was measured by flow cytometry for GFP at day 1 in samples treated with AAV6 but not ZFN mRNA. (C-D) Cell viability (C) and total cell counts by hemocytometer (D) were measured one day after genome editing. Conditions were pooled across AAV6 MOIs. Data are shown as mean ± SEM. *** *p* < 0.001.

## Discussion

AAV vectors are invaluable reagents for site-specific genome editing of human hematopoietic cells, with AAV6 serotypes in particular being widely used to deliver homology donors to HSPCs,^2, 4, 11, 12^ T cells,^13–15^ and B cells.^16, 17, 19^ The *in vitro* tropism of AAV6 for human hematopoietic cells,^2, 13, 16^ as well its weak induction of innate immune pathways that could trigger harmful biological consequences in engineered cells,^41, 42^ may contribute to its success. In addition, the vector’s ability to transduce both dividing and quiescent cells^43^ may also be beneficial, particularly if the genome is used as a homology template only after second-strand synthesis as some have suggested,^7^ though the G2/S phase restriction of cellular factors required for HDR^25^ may limit the advantages conferred by this attribute.

Despite using similar protocols, we and others have observed relative inefficiencies in AAV6 transduction and genome editing for human lymphocytes compared to HSPCs. In this study we chose to focus on optimizing culture conditions for AAV6 transduction of suspension cells *ex vivo*, hypothesizing that achieving efficient delivery of the homology template may be (one of) the rate-limiting steps in these experiments.

Our initial experiments focused on the impact of serum on AAV transduction, based on reports that FBS could be inhibitory.^13^ We observed dose-dependent inhibition of AAV6 transduction with both fetal bovine and human serum, which functioned by blocking AAV6 attachment to target cells. Inhibition by serum varied across different lots of FBS and AAV serotypes and was observed despite differing use of glycan receptors between the serotypes tested (AAV1: sialic acid, AAV2: heparan sulfate proteoglycan, AAV6: both).^28^ This variance in neutralization across lots and AAV serotypes is consistent with the presence of anti-AAV neutralizing antibodies (NABs). NABs to therapeutically relevant AAV serotypes, including AAV6, have been found in a variety of animal models, including nonhuman primates, rodents, dogs, sheep, cats, horses, and pigs.^44–51^ However, we are not aware of any studies that have reported the presence of anti-AAV NABs in FBS.

We also investigated the impact of other culture conditions on AAV6 transduction. In line with previous reports,^37^ we found that increasing cell density by reducing the volume of media could greatly enhance AAV6 transduction. We were able to culture 10^6^ primary hematopoietic cells in volumes as small as 10 μL using 96-well U-bottom plates (10^8^ cells/mL, ~100-fold greater than a standard culture concentration), with minimal impact on cell viability during a 1 h transduction. This short incubation period matched our observations in K562 cells that most of the benefits of AAV6 transduction at high densities are realized during the first hour, in an initial logarithmic phase that then tapers off to further increase at a linear rate.

The biphasic pattern of AAV6 transduction that we observed could represent an initial excess of molecules involved in AAV6 attachment or entry, allowing a rapid initial burst of transduction, followed by a slower, linear phase that could be dependent on recycling or *de novo* synthesis of these receptors. Accordingly, previous studies using AAV2 and adherent cells have suggested that endocytosis may be a rate-limiting step in AAV transduction.^52, 53^ However, the reported kinetics of AAV uptake have varied,^36, 52, 54, 55^ perhaps reflecting differences in detection methods (physical labeling of particles vs. productive gene expression), AAV serotype, or in endocytic pathway usage. Several different endocytic pathways have all been implicated in AAV uptake, and usage may vary across cell types.^56, 57^ The unique biphasic transduction kinetics revealed here by an extended AAV6 transduction timeline in K562 cells may warrant further study to confirm these findings and elucidate the mechanisms involved, which could lead to novel methods to modulate the rate of AAV endocytosis in cells.

The importance of endocytosis in this system is also reflected in the ability of electroporation to increase AAV transduction and editing rates, independent of the presence of a targeted nuclease. Charlesworth *et al*. have previously suggested that electroporation prior to AAV6 transduction can prime cells to increase endocytosis, allowing greater AAV6 uptake through AAVR-mediated pathways.^21^ Similarly, we observed that electroporation enhanced AAV6 transduction of K562 cells, with the additional finding that this enhancement occurred regardless of whether AAV6 was added to cells before or after electroporation. Moreover, when cells were exposed to AAV6 prior to electroporation, transduction was dependent on viral attachment. The lack of effect of washing unbound AAV6 out prior to electroporation suggests enhanced entry of viral particles already attached to the cell rather than, for example, a bulk effect on direct membrane permeation caused by electroporation.

Based on the findings in K562 cells, we designed an optimized protocol for AAV6 transduction of hematopoietic cells that comprised a 1-hr incubation in concentrated cell culture conditions in serum-free media, followed by electroporation with ZFN mRNA, and then a switch to culture in serum-containing media immediately afterwards. We believe this timing maximizes AAV6 transduction while minimizing the potential negative impacts of serum starvation or cell overcrowding. Since our results here suggest that electroporation can enhance AAV transduction even after cells are exposed to the vector, and we have previously shown that this sequence of events is compatible with site-specific genome editing,^2^ we chose to transduce the cells prior to electroporation. In this way, the high-density culture is performed when the cells are fully healthy rather than recovering from the harsher electroporation procedure.

We compared this optimized protocol to typical protocols described in the literature,^13, 14, 16^, with the addition that the AAV6 transduction step was always performed in the absence of FBS. In both primary human T and B cells, we observed significant advantages with the optimized protocol compared to an overnight serum-free AAV transduction following electroporation, both in cell viability and genome editing efficiency. Editing rates were improved by up to 7-fold in T cells and 21-fold in B cells at equal MOIs, and similar frequencies were achieved with 50-100-fold less AAV6 per cell. These improvements could significantly reduce the amount of AAV6 required for *ex vivo* genome editing of lymphocytes, which could represent a significant cost savings. This is especially apparent in B cells, where the difficulty in transducing these cells with AAV6 means that MOIs of 10^6^ are required without the optimized high-density transduction. At this MOI, editing 1 million cells *ex vivo* would require 10^12^ vg, which is 10-100 times more AAV than some therapeutic doses for *in vivo* hepatic gene transfer in mice.^58^

Our data also highlight the impact of factors beyond AAV6 transduction on the efficiency of HDR-mediated genome editing, since an upper limit of editing rates clearly existed that could not be surpassed by increasing the AAV6 MOI. Others factors involved include the rate of DSB induction by the targeted nuclease, the influence of local DNA sequence on the choice of DNA repair pathway used,^59^ and the activity of HDR repair pathways as a function of the cell cycle.^25^ This last point may be of particular relevance as the overnight serum starvation in the original protocols may have caused significant growth arrest and exit from the cell cycle into a G0 phase^40^ that does not support HDR. This could limit the ability to convert AAV6 transduction into site-specific genome editing, and may at least partially explain the advantage of the optimized protocol that does not involve prolonged serum deprivation. Superior serum-free media formulations may be better able to support cell growth while avoiding serum-mediated inhibition of AAV transduction.

Indeed, the use of more effective serum-free media in HSPCs is one hypothesis for why the optimized protocol was unable to enhance genome editing in these cells, in contrast to findings in CD4^+^ T cells and B cells. Although serum-containing media was used after genome editing in HSPCs for historical reasons and to maintain similarities between the cell types studied in this manuscript, we and others have found that the serum-free media used here can efficiently support growth of HSPCs without additional FBS supplementation (not shown).^11^ Thus, cells may have been healthier after the original protocol and better able to undergo HDR-mediated genome editing, reducing a potential advantage of the optimized protocol. Alternatively, the lack of improvement with the optimized protocol in HSPCs may reflect the much greater permissivity of these cells to AAV6 transduction. Compared with MOIs of 5 x 10^4^ in T cells and 10^6^ in B cells, only 3 x 10^3^ vg/cell were originally required for efficient genome editing in HSPCs. Our findings in K562 cells suggest that the impact of reduced media volumes is less dramatic at lower AAV MOIs across the conditions tested, perhaps suggesting that significantly lower volumes would be required to see an improvement in AAV transduction at lower MOIs in HSPCs.

In summary, we empirically tested cell culture parameters for their impact on transduction of hematopoietic cells by AAV6 vectors, and used these findings to design an optimized protocol for genome editing. In primary human T and B lymphocytes, this approach significantly improved site-specific genome editing in conjunction with electroporation of ZFN mRNA, and we have also observed similar effects when using electroporation of CRISPR/Cas9 RNPs (data not shown). The dramatic differences we observed across protocols suggest that more detailed reports of cell culture and AAV transduction methodology may be necessary to ensure good reproducibility across labs. Our results also highlight the importance of cell-intrinsic factors and optimizing growth conditions to enable the highest rates of *ex vivo* genome editing, and suggest that improved serum-free media may be required for continuing enhancements in *ex vivo* genome editing. Nevertheless, these procedures may be a useful starting point for investigators using AAV6 for site-specific genome editing in hematopoietic cells.

## Materials and Methods

### AAV vectors

AAV6-CCR5-GFP vectors containing AAV2 ITRs, CCR5 homology arms of 473 bp (left) and 1431 bp (right), a hPGK promoter driving eGFP, and a BGH polyA signal were produced as previously described^2^ and generously provided by Sangamo Therapeutics. AAV1-CMV-GFP and AAV2-CMV-GFP vectors were purchased from Vigene Biosciences (Rockville, MD). AAV vectors were titrated as previously described,^39^ and protocols are provided in the Supplemental Information.

### Cell line culture, AAV transduction, and electroporation

HEK-293T cells and HeLa cells were cultured in DMEM supplemented with 10% FBS and 1% penicillin/streptomycin. Cells were seeded overnight to adhere to plates and washed once with PBS prior to AAV transduction in DMEM, with or without FBS or human AB serum. FBS was heat-inactivated at 56°C water bath for 30 mins. AAV vectors were added to cells at indicated MOIs, and after 4 h at 37°C, 10% FBS (final volume) was restored to the culture if appropriate.

K562, Raji, and Molt4.8 cells were cultured in RPMI-1640 medium supplemented with 10% FBS and 1% penicillin/streptomycin. Cells were washed twice with PBS, seeded into plates at indicated cell concentrations in RPMI-1640, with or without FBS or human AB serum, and transduced with AAV vectors at indicated MOIs at 37°C. After 2 h (or as indicated), 10% FBS (final volume) was restored to the culture if appropriate. Electroporation of K562 cells was performed using a SF Cell Line 4D-Nucleofector kit and 4D-X Nucleofector using pulse code FF-120 (Lonza, Basel, Switzerland), per the manufacturer’s recommendations. After 2 days, GFP expression was measured by flow cytometry and vector copy numbers were measured by ddPCR as previously described.^39^ Detailed protocols for AAV copy number determination are provided in the Supplemental Information.

### Viral attachment assay

K562 cells were washed with PBS, resuspended in RPMI-1640 with or without 10% FBS, and incubated at 4°C for 30 mins at 10^6^ cells/mL. AAV6-CCR5-GFP vectors were added at an MOI of 10^4^ and allowed to attach to the cells for 1 h at 4°C. Cells were washed at 4°C to remove unattached AAV6 virions, resuspended in RPMI-1640 with or without 10% FBS as indicated, and incubated at 37°C. Control samples were also transduced with AAV6-CCR5-GFP in FBS-free RPMI-1640 at the same MOI and cell concentration. After allowing transduction for 2 h, samples without FBS were supplemented with 10% FBS, and cells were cultured for 2 days at 37°C.

### Human CD4^+^ T cells

Human buffy coat preparations were obtained from Gulf Coast Regional Blood Center (Houston, TX). PBMCs were isolated by Ficoll-Paque (GE Healthcare Life Sciences, Marlborough, MA) density centrifugation using Leucosep tubes (Greiner Bio-One, Kremsmünster, Austria) and platelets were reduced by low speed washing. Human CD4^+^ T cells were isolated by positive magnetic selection using human CD4 MicroBeads kit (Miltenyi Biotec, Bergisch Gladbach, Germany) per manufacturer’s instructions. For activation, purified CD4^+^ T cells were cultured at 2 x 10^6^ cells/mL in T cell media: X-VIVO-15 media supplemented with 10% FBS, 2 mM L-glutamine, 1% penicillin/streptomycin/amphotericin B (Sigma-Aldrich), 20 ng/mL IL-2 (Peprotech, Rocky Hill, NJ), and Dynabeads Human T-Activator CD3/CD28 (Thermo Fisher) at 1 bead per cell, as previously described.^13^ After 3 days, beads were removed using a DynaMag-2 (Thermo Fisher) per manufacturer’s instructions. Cells were washed twice with PBS, then genome edited with indicated protocols.

### Human B cells

Frozen human peripheral blood CD19+ B cells were purchased from StemCell Technologies (Vancouver, Canada). Cells were thawed per manufacturer’s instructions and cultured as previously described.^60^ Briefly, cells were initially activated at 4 x 10^5^ - 10^6^ cells/mL in B cell activation media: Iscove’s Modified Dulbecco’s Medium (IMDM, Corning Inc., Corning, NY) supplemented with 10% FBS, 5 μg/mL soluble CD40L (R&D Systems, Minneapolis, MN), 10 μg/mL anti-His tag antibody (clone AD1.1.10, R&D Systems), 50 ng/mL CpG ODN 2006 (Invivogen, San Diego, CA), 10 ng/mL IL-2 (R&D Systems), 50 ng/mL IL-10, and 10 ng/mL IL-15 (Peprotech). After 2 days of activation, cells were washed twice with PBS, then genome edited with indicated protocols.

After genome editing, cells were cultured an additional 2 days in B cell activation media. Then, cells were pelleted by centrifugation and media was replaced with plasmablast generation media: IMDM supplemented with 10% FBS, 10 ng/mL IL-2 (R&D Systems), 50 ng/mL IL-6 (Peprotech), 50 ng/mL IL-10, and 10 ng/mL IL-15. After a further 3 days of culture, cells were pelleted by centrifugation and media was replaced with plasma cell generation media: IMDM supplemented with 10% FBS, 50 ng/mL IL-6, 10 ng/mL IL-15, and 500 U/mL IFN-α (R&D Systems). Cells were cultured in this media for 3 additional days.

### Human CD34^+^ HSPCs

Fetal liver CD34^+^ HSPCs were isolated from tissue obtained from Advanced Bioscience Resources (Alameda, CA) as anonymous waste samples, with approval of the University of Southern California’s Institutional Review Board. CD34^+^ cells were isolated as previously described,^2^ using physical disruption, incubation in collagenase to give single cell suspensions, and magnetic bead selection using an EasySep™ Human CD34 Positive Selection Kit (STEMCELL Technologies Inc.). The resulting CD34^+^ HSPCs were cultured in HSPC media: StemSpan SFEM II (STEMCELL Technologies Inc.) supplemented with 1% penicillin/streptomycin/amphotericin B and SFT cytokines: 50 ng/mL each of SCF, Flt3 ligand and TPO (R&D Systems). After overnight pre-stimulation, HSPCs were washed twice with PBS, then genome edited with indicated protocols.

### Genome editing protocols

Protocols for preparation of ZFN reagents, measurement of indels, and detailed protocols for genome editing of K562 cells, human CD4 T cells, human B cells and human HSPCs are provided in the Supplemental Information.

### Flow cytometry

GFP expression was measured by flow cytometry as indicated on either a FACSCanto II (BD Biosciences, San Diego, CA) or Guava easyCyte (MilliporeSigma, Burlington, MA). Viability was measured by 7-AAD staining (BD Biosciences). Data were analyzed using FlowJo software (Flowjo LLC, Ashland, OR).

### Statistics

Results are reported as means ± standard error of the mean (SEM). Significant differences between groups were determined with unpaired Student’s t test, one-way analysis of variance with Tukey posttests, two-way analysis of variance with Bonferroni posttests, linear regression, or semilogarithmic regression, as appropriate. *p* values of < 0.05 were considered significant. Analyses were performed using GraphPad Prism software (San Diego, CA). Differences are indicated as * *p* < 0.05, ** *p* < 0.01, *** *p* < 0.001, and ns = not significant.

## Acknowledgements

We would like to thank B. Riley, M. Holmes, and Sangamo Therapeutics Inc. for providing AAV6-CCR5-GFP vectors, CCR5 ZFN reagents, and discussions about methods for B cell culture and genome editing. This work was supported by National Institutes of Health grants HL129902 and HL156274 to P.M.C. G.L.R. was supported by a Career Development Award from the American Society of Gene & Cell Therapy. The content is solely the responsibility of the authors and does not necessarily represent the official views of the American Society of Gene & Cell Therapy. H-Y.C. was supported by a Taiwan USC scholarship.

## Author contributions

G.L.R., C.H., and R.C. performed experiments. G.L.R. and P.M.C. designed experiments and analyzed and interpreted data. E.S. and H-Y.C. contributed to discussions. G.L.R. and P.M.C. wrote the manuscript. P.M.C. supervised the study.

