## Supplemental Information for "Optimization of AAV6 transduction enhances site-specific genome editing of primary human lymphocytes"

**Supplemental Figures**

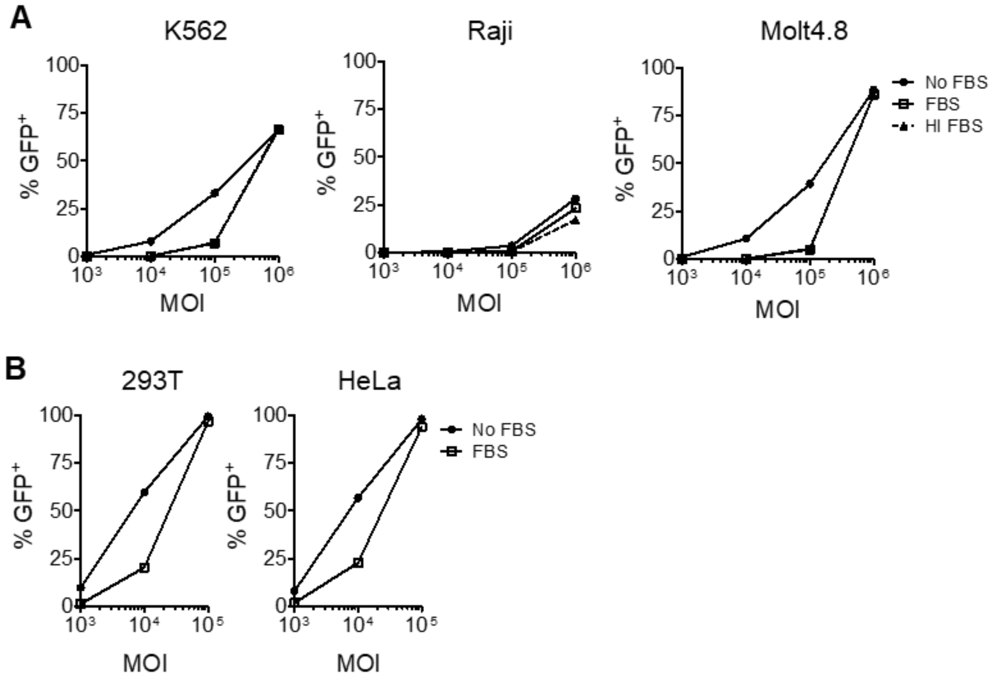


**Figure S1. Raw data for inhibition of AAV6 cell line transduction by FBS.** GFP expression measured at day 2 by flow cytometry is shown for transduction with AAV6-CCR5-GFP vectors at indicated MOIs with or without FBS for (A) suspension K562, Raji, and Molt4.8 cells, or (B) adherent 293T and HeLa cells. Percent inhibition in Fig. 1A-B and Fig. S2 were calculated by comparing these data with and without FBS. Associated with Fig. 1.

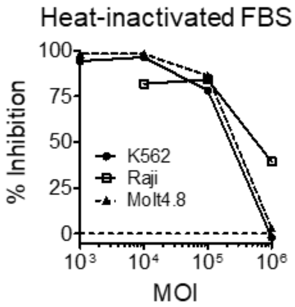


**Figure S2. Inhibition of AAV6 transduction by heat-inactivated FBS.** Percent inhibition of AAV6-CCR5-GFP transduction on suspension cell lines (K562, Raji, and Molt4.8) by 10% FBS was calculated across the indicated MOIs from GFP expression at day 2 by flow cytometry. Associated with Fig. 1.

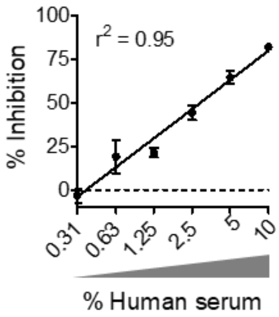


**Figure S3. Dose-dependent inhibition of AAV6 transduction by human serum.** Percent inhibition of AAV6-CCR5-GFP transduction (MOI = $10^4$) was calculated in K562 cells over a range of human serum concentrations, and a semi-logarithmic regression line was calculated as shown. Data are shown as mean ± SEM for $n$ = 3 technical replicates. Associated with Fig. 1.

| MOI | $10^3$ | $3.16 \times 10^3$ | $10^4$ | $3.16 \times 10^4$ | $10^5$ | $3.16 \times 10^5$ |
|---|---|---|---|---|---|---|
| Slope | -0.91±0.33 | -3.31±0.38 | -11.42±0.88 | -21.3±2.31 | -32.23±1.72 | -19.9±1.86 |
| Y-intercept | 3.16±0.41 | 9.15±0.59 | 32.2±1.62 | 62.2±4.68 | 106.6±3.85 | 93.6±4.57 |
| $R^2$ | 0.5225 | 0.8820 | 0.9282 | 0.8950 | 0.9804 | 0.9665 |

**Table S1. Linear regression characteristics for the impact of cell culture volume and MOI on AAV6 transduction.** Associated with Fig. 2A.

| **MOI** | **$10^4$** | **$10^4$** | **$10^3$** |
|---|---|---|---|
| **Cells/mL** | **$10^6$** | **$10^7$** | **$10^7$** |
| Slope | 1.53 ± 0.04 | 1.57 ± 0.15 | 0.77 ± 0.04*** |
| Y-intercept | 16.14 ± 0.50 | 32.9 ± 2.01*** | 5.99 ± 0.60 |
| $R^2$ | 0.98 | 0.90 | 0.96 |

**Table S2. Linear regression characteristics for AAV6 transduction over time.** Associated with Fig. 2C-D. *** $p < 0.0001$.

### Supplemental Methods

#### AAV vector titration

All AAV vectors were titrated as previously described.[1] Briefly, DNA was extracted by treatment with DNaseI (New England Biolabs, Ipswich, MA) to remove residual plasmid DNA, followed by treatment with proteinase K (Sigma-Aldrich, St. Louis, MO) for 1 hour at 37°C. The extracted DNA was stored at -20°C until titration. AAV vector genome (vg) titers were determined by TaqMan qPCR (Thermo Fisher, Waltham, MA) using ITR specific primers (AAV ITR-Forward 5'-GAACCCCTAGTGATGGAGTT-3', AAV ITR-Reverse 5'-CGGCCTCAGTGAGCGA-3') and probe (AAV ITR-Probe 5'-FAM-CACTCCCTCTCTGCGCGCTCG-Tamra-3'). To prepare the standard curve, serial dilutions of DNA extracted contemporaneously from a recombinant AAV2 Reference Standard Material (American Type Culture Collection; VR-1616) was used.[2]

#### AAV vector copy number analysis

Cells were pelleted by centrifugation and DNA was isolated using the DNeasy Blood & Tissue Kit (Qiagen). Roughly 5 ng of genomic DNA was mixed with primers and probes for a human RPP30 copy number assay labeled with HEX (Bio-Rad, Hercules, CA) as an internal control, and GFP-specific primers and probe were as follows: Forward 5'-AGCAAAGACCCCAACGAGAA-3', Reverse 5'-GGCGGCGGTCACGAA-3', Probe 5'-FAM-CGCGATCACATGGTCCTGCTGG-3'. Droplets were prepared using ddPCR Supermix for Probes (No dUTP) and a QX200 Droplet Generator (Bio-Rad). The PCR reaction was run on a C1000 Touch Thermal Cycler with the following conditions: 95 °C 10 min, 40 cycles (94° C 30 s, 60° C 1 min), 98° C 10 min, and 4° C forever. After the PCR reaction, the samples were read on a QX200 Droplet Reader and the data were analyzed with QuantaSoft analysis software (Bio-Rad). Linear regression analysis was performed using GraphPad Prism software (San Diego, CA).

#### ZFN reagents and mRNA production

ZFNs targeting the CCR5 locus using a bicistronic cassette with a 2A peptide have been described previously.[1, 3] Plasmid DNA was linearized by restriction enzyme digest (SpeI) and purified using the Zymo DNA Clean and Concentrator protocol (Zymo Research, Irvine, CA). *In vitro* transcription of mRNA was performed using the T7 mScript Standard mRNA Production System (Cellscript, Madison, WI) per manufacturer's instructions. RNA was purified using RNA Clean & Concentrator-25 (Zymo Research) per manufacturer's protocol and stored at -80° C until use.

#### Detection of indels

Cells were pelleted and genomic DNA was extracted using a NucleoSpin Tissue kit (Takara Bio Inc., Kusatsu, Shiga, Japan) per manufacturer's instructions. The human CCR5 locus was amplified with AmpliTaq Gold 360 Master Mix (Thermo Fisher) with the following primers: Forward 5'-AAGATGGATTATCAAGTGTCAAGTCC-3', Reverse 5'- CAAAGTCCCACTGGGCG-3'. PCR products were then digested using the GeneArt Genomic Cleavage Detection Kit (Thermo Fisher), and DNA was visualized on a Mini-PROTEAN 5% TBE gel (Bio-Rad) stained with GelRed Nucleic Acid Stain (Biotium, Fremont, CA).

#### Detailed protocols for K562 cell genome editing

Genome editing protocols in K562 cells were as follows. Original protocol: K562 cells were washed twice with PBS, then 2 x $10^5$ K562 cells were Nucleofected in 20 µL SF buffer, as above, along with 4 µg CCR5-specific ZFN mRNA. Cells were resuspended in serum-free RPMI-1640 and transduced with AAV6-CCR5-GFP at indicated MOIs for 2 h before media was supplemented with 10% FBS. Optimized protocol: K562 cells were washed twice with PBS, then resuspended at indicated concentrations in 10-20 µL serum-free RPMI-1640. Cells were transduced with AAV6-CCR5-GFP at indicated MOIs for the indicated time, then mixed with 80-90 µL buffer SF and Nucleofected as above with ZFN mRNA. In both cases cells were cultured in RPMI-1640 supplemented with 10% FBS for 3 weeks to measure stable GFP gene insertion by flow cytometry.

#### Detailed protocols for human $CD4^+$ T cell genome editing

For the original protocol: $10^6$ cells were resuspended in 100 µL BTXpress Electroporation Buffer (Harvard Apparatus, Holliston, MA), mixed with 8 µg ZFN mRNA, and electroporated using a BTX ECM 830 (Harvard Apparatus) at 250 V for 5 ms. Cells were diluted to $10^6$ cells/mL in serum-free T cell media supplemented with 20 ng/mL IL-7 (Peprotech) and transduced with AAV6-CCR5-GFP homology donor vectors at indicated MOIs at 37°C. After overnight incubation (16-24 h), media was supplemented with 10% FBS and cells were cultured for 10 days to measure stable GFP genome editing by flow cytometry.

For the optimized protocol: $10^6$ cells were resuspended in 10 µL serum-free T cell media and transduced with AAV6-CCR5-GFP homology donor vectors, at indicated MOIs, in 96-well U-bottom plates for 1 h at 37°C. Then, cells were mixed with 90 µL BTXpress Electroporation Buffer and 8 µg ZFN mRNA, and electroporated using a BTX ECM 830 at 250 V for 5 ms. After electroporation, cells were diluted to $10^6$ cells/mL in T cell media containing FBS supplemented with 20 ng/mL IL-7. Cells were cultured for 10 days to measure stable GFP genome editing by flow cytometry.

**Detailed protocols for human B cell genome editing**

For the original protocol: cells were resuspended at 2 x $10^6$ cells/mL in BTXpress Electroporation Buffer, mixed with 8 µg ZFN mRNA, and electroporated using a BTX ECM 830 at 250 V for 5 ms. Cells were then resuspended at 4 x $10^5$ cells/mL in serum-free B cell activation media and transduced with AAV6-CCR5-GFP homology donor vectors at indicated MOIs overnight (16-24 h) at 37°C. The next day, media was supplemented with 10% FBS and further culturing/differentiation was performed.

For the optimized protocol: cells were resuspended at 2-5 x $10^7$ cells/mL in 10 µL of serum-free B cell activation media and transduced with AAV6-CCR5-GFP homology donor vectors at indicated MOIs in 96-well U-bottom plates for 1 h at 37°C. Then, cells were mixed with 90 µL BTXpress Electroporation Buffer and 8 µg ZFN mRNA, and electroporated using a BTX ECM 830 at 250 V for 5 ms. After electroporation, cells were diluted to 4 x $10^5$ - 1 x $10^6$ cells/mL in B cell activation media with serum and further culturing/differentiation was performed.

**Detailed protocols for human CD34$^+$ HSPC genome editing**

For the original protocol: $10^6$ cells were resuspended in 100 µL BTXpress Electroporation Buffer, mixed with 8 µg ZFN mRNA, and electroporated using a BTX ECM 830 at 250 V for 5 ms. Cells were resuspended in serum-free HSPC media and transduced with transduced with AAV6-CCR5-GFP homology donor vectors at indicated MOIs for 4 h at 37°C. After 4 h, media was supplemented with 10% FBS and cells were cultured for 10 days to measure stable GFP genome editing by flow cytometry.

For the optimized protocol: $10^6$ cells were resuspended in 10 µL serum-free HSPC media and transduced with AAV6 vectors at indicated MOIs in 96-well U-bottom plates for 1 h at 37°C. Then, cells were mixed with 90 µL BTXpress Electroporation Buffer and 8 µg ZFN mRNA, and electroporated using a BTX ECM 830 at 250 V for 5 ms. After electroporation, cells were diluted to $10^6$ cells/mL in HSPC media supplemented with 10% FBS and cultured for 10 days to measure stable GFP genome editing by flow cytometry.
